# Treatment with anti-HER2 chimeric antigen receptor tumor-infiltrating lymphocytes (CAR-TILs) is safe and associated with antitumor efficacy in mice and companion dogs

**DOI:** 10.1101/2022.09.11.507449

**Authors:** Elin MV Forsberg, Rebecca Riise, Sara Saellström, Joakim Karlsson, Samuel Alsén, Valentina Bucher, Akseli Hemminki, Roger Olofsson Bagge, Lars Ny, Lisa M Nilsson, Henrik Rönnberg, Jonas A Nilsson

## Abstract

Patients with metastatic melanoma have a historically poor prognosis, but recent advances in treatment options, including targeted therapy and immunotherapy, have drastically improved the outcomes for some of these patients. However, not all patients respond to available treatments, and around 50% of patients with metastatic cutaneous melanoma and almost all patients with metastases of uveal melanoma die of their disease. Thus, there is a need for novel treatment strategies for patients with melanoma that do not benefit from the available therapies. Chimeric antigen receptor-expressing T (CAR-T) cells are largely unexplored in melanoma. Traditionally, CAR-T cells have been produced by transducing blood-derived T cells with a virus expressing CAR. However, tumor-infiltrating lymphocytes (TILs) can also be engineered to express CAR, and such CAR-TILs could be dual-targeting. To this end, tumor samples and autologous TILs from metastasized human uveal and cutaneous melanoma were expanded *in vitro* and transduced with a lentiviral vector encoding an anti-HER2 CAR construct. When infused into patient-derived xenograft (PDX) mouse models carrying autologous tumors, CAR-TILs were able to eradicate melanoma, even in the absence of antigen presentation by HLA. To advance this concept to the clinic and assess its safety in an immune-competent and human-patient-like setting, we treated four companion dogs with autologous anti-HER2 CAR-TILs. We found that these cells were tolerable and showed signs of anti-tumor activity. Taken together, CAR-TIL therapy is a promising avenue for broadening the tumor-targeting capacity of TILs in patients with checkpoint immunotherapy-resistant melanoma.

## Introduction

Patients with metastatic melanoma have a historically poor prognosis; however(1), recent advances in treatment options have drastically improved patient prognosis. Targeted therapies using inhibitors of BRAF alone (2,3) or in combination with MEK inhibitors (4,5) have shown good response rates in patients with metastatic cutaneous melanoma. However, most patients treated with these inhibitors develop drug resistance. Immunotherapies, including checkpoint inhibitors targeting PD1 and CTLA4 or LAG3, can result in more durable response rates among patients with melanoma(6–9). Adoptive T cell transfer (ACT) with tumor-infiltrating lymphocytes (TILs) has also been used to treat metastatic melanoma in clinical trials, with response rates of approximately 50%(10–12). Importantly, not all patients with metastatic malignant melanoma respond to current treatment strategies; therefore, alternative and/or combination therapies are currently being explored in preclinical experiments and trials.

Uveal melanoma (UM) is a rare form of melanoma (13) arising in the uveal tract of the eye, i.e. the iris, ciliary body and choroid. UM is treated with brachytherapy or enucleation with very good local control (97%) (14). However, approximately 50% of patients will later present with metastatic disease(15), mainly to the liver, but also to other sites (16). Patients with spread UM rarely respond to systemic chemotherapy or targeted therapies, (17) and combined immune checkpoint inhibitors have not shown the same promising effect in UM (18,19) as in cutaneous melanoma (20). Benchmark data suggest an average progression-free and overall survival of approximately 3.3 and 10.2 months, respectively (21). Loco-regional treatment with isolated hepatic perfusion or percutaneous hepatic perfusion demonstrates high response rates and prolonged progression-free survival in randomized trials, but mature survival data are still pending (22,23). The combined treatment with PD-1 inhibitor pembrolizumab and the HDAC inhibitor entinostat exhibited durable responses in a fraction of patients, including one with an iris melanoma with a high mutation burden and patients with tumors exhibiting a wildtype *BAP1* tumor suppressor gene (24). ACT with TILs was tested in patients with UM in a clinical trial, with a response rate of 35% (25). Finally, for patients with the HLA-A2 genotype, the bispecific T cell engager tebentafusp can activate anti-tumor immune responses (26) and prolong the survival of patients with UM (27), despite surprisingly low response rates (28). Hence, although recent progress has been made, metastatic UM remains a medical challenge.

Immunotherapies aim to overcome tumor immune evasion strategies to re-activate the patient’s own immune system to attack and kill the tumor. However, some tumors downregulate the antigen presentation pathway (29,30), rendering these tumor cells insensitive to TCR-mediated recognition and killing, thereby disarming both immune checkpoint inhibition and ACT. One way to overcome this problem is to equip T cells with a chimeric antigen receptor (CAR) designed to recognize a specific protein expressed on the surface of tumor cells irrespective of antigen presentation. CAR-T therapy has not yet been approved for use in any solid cancer; however, CD19 CAR-T therapy is used in young patients with acute lymphocytic leukemia (ALL) (31). The reason why CAR-T therapy has not yet been successful in solid cancers is not fully understood but includes heterogeneous expression of antigens, expression of checkpoint proteins, poor homing and tissue penetrance of CAR-T cells, an immune suppressive tumor microenvironment (TME), and CAR-T cell endurance (32).

Mouse models are useful for studying the antitumor efficacy of human CAR-T cells, but there are limitations. Patient-derived xenograft (PDX) models are immunodeficient (33), and in both PDX models and cell line-derived xenograft models, the tumor stroma and off-target tissues of CAR-T cells are of mouse origin. Therefore, these models cannot be readily used for TME or toxicity studies without additional genetic engineering. Neither xenograft nor syngeneic transplant models are spontaneous tumor models; therefore, tumor architecture can also be suboptimal. Companion dogs are an emerging and complementary model to study toxicity and anti-cancer treatment efficacy (34). Dogs live and socialize with their human owners, share their habits and microbiota, and develop lifestyle diseases, such as cardiovascular problems, joint problems, diabetes, and cancer, with age (35). Malignancies range from leukemia and lymphoma to solid tumors such as mammary or squamous cell carcinoma of the head and neck (36). Melanoma is a particularly aggressive form of cancer in some breeds of dogs (37). It is not associated with solar damage and most often develops in mucosal areas such as the mouth or under the nail bed. Similar to human mucosal melanoma, the spectrum and driver mutations of canine melanoma are different from those of human cutaneous and uveal melanoma (38,39).

We previously reported good antitumor efficacy of CAR-T cells directed against HER2 in PDX models of cutaneous and uveal melanoma (40). We demonstrated that the elicited effect of CAR-T cells was target-specific, as CRISPR/Cas9-mediated disruption of HER2 abolished the sensitivity of melanoma cells to anti-HER2 CAR-T cells. Importantly, CAR-T cells were also able to eradicate tumors that were refractory to ACT. Current CAR-T therapies use CAR-transduced T-cells from blood as drug substances. This T cell pool largely consists of naïve and memory T cells carrying a TCR with irrelevant affinity. Naïve T cells generally do not express molecules that facilitate homing to inflamed peripheral tissues. TILs, on the other hand, can home to tumors; therefore (41,42), they could potentially serve as good starting materials for the generation of CAR-T cells. The aim of this study was to assess whether CAR expression in TILs can boost tumor cell death. We also assessed whether the CAR-TILs were safe and tolerable in mice and companion dogs.

## Results

### TILs differentially express chemokine receptors and selectins

CAR-T cells have been approved for clinical use against some B-cell malignancies; however, no CAR-T cell therapy has been approved for solid tumors. To investigate the difference in the expression of cell surface molecules involved in T cell trafficking and homing, we performed single-cell sequencing of blood-derived T cells and young TILs that had grown out uveal melanoma metastases in the presence of the T cell growth factor interleukin-2 (IL-2) (43). Focusing on differentially expressed genes in the category of ‘chemokine receptors’ or ‘adhesion molecules’ we find that genes like *ITGA4* (CD49D), *ITGB2* (CD18), *ITGAL* (CD11a), *ITGAE* (CD103), *PECAM1* (CD31) and *CXCR3* are all more highly expressed on TILs compared to on blood-derived T cells (**Suppl Fig S1A**). This observation was validated at the protein level using flow cytometry, which demonstrated that at least some of this differential expression could be caused by the IL-2 used in the culture medium of TILs (**Suppl Fig S1B-C**).

### CAR-TILs can kill melanoma cells in vitro

CAR-T cell manufacturing is performed by modifying blood-derived T cells from leukophoresis with a chimeric antigen receptor (CAR) that binds to a surface antigen on the tumor cell. To investigate whether anti-HER2 CAR-expressing TILs, hereafter called CAR-TILs, are capable of recognizing melanoma cells, we used TILs from human patients with melanoma and transfected these cells with an mRNA encoding a HER2 CAR consisting of a single-chain variable fragment (scFv) to detect HER2 fused with the signaling domains of CD3 epsilon and the CD28 co-stimulatory molecule (40). We consistently observed HER2 CAR expression in these cells, as assessed by HER2 binding to CAR-TILs (**Fig 1A** and **Suppl Fig S2**). CAR was functional and specific because the CAR-TILs (but not mock-transfected TILs) from patient MM1 degranulated, released, and caused accumulation of interferon gamma (IFN-γ) in the medium when co-cultured with the HER2 positive uveal melanoma cell line 92-1. This was dependent on HER2 expression since CRISPR-generated 92-1 HER2 knockout cells (40) were not able to activate CAR-TILs. This demonstrates that anti-HER2 CAR-TILs can react with HER2 on the surface of melanoma cells.

**Figure 1.**
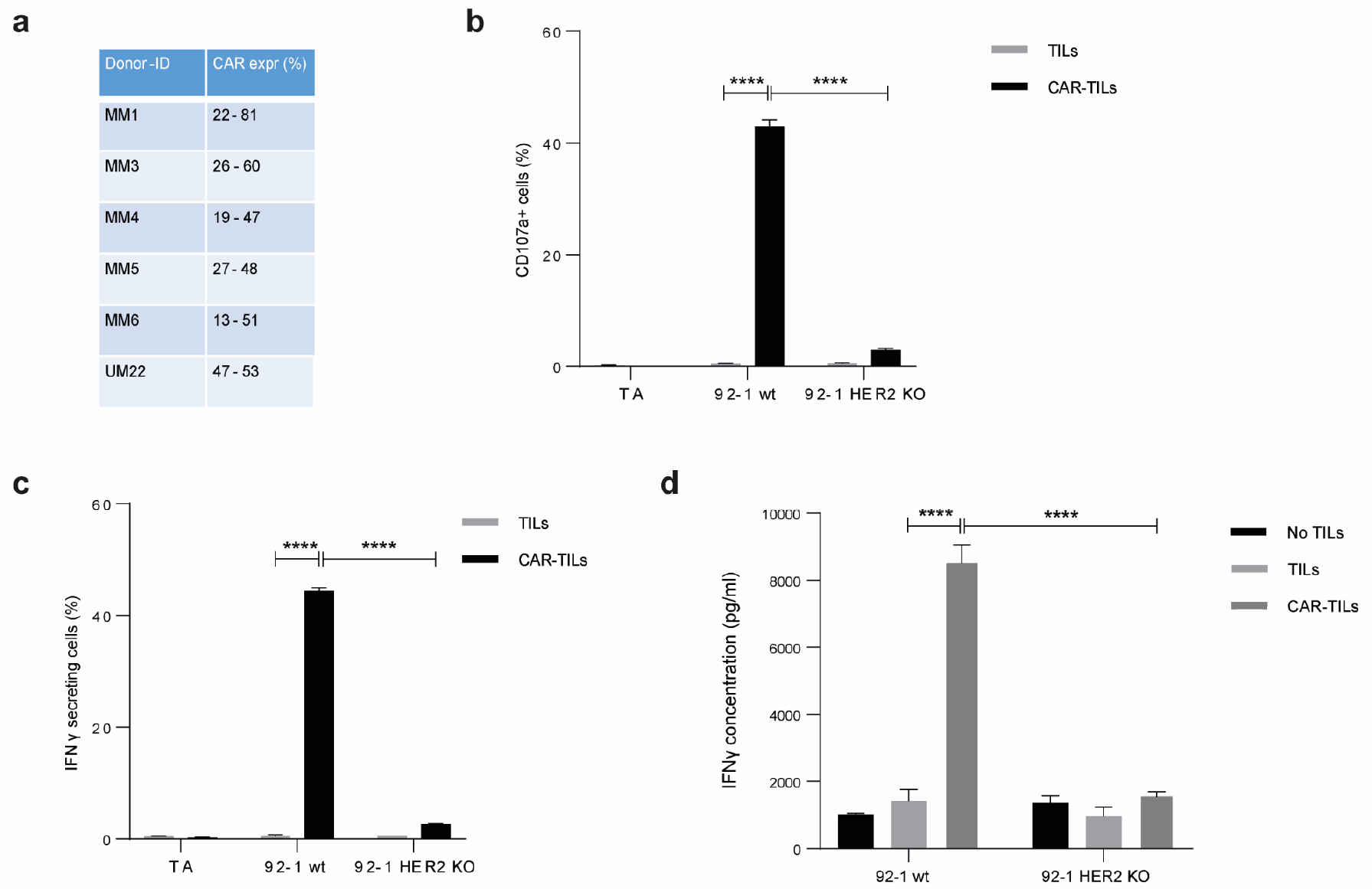
TILs can be activated by equipping them with a CAR construct. mRNA encoding anti-HER2 CAR was electroporated into TILs from five cutaneous melanomas (MM1, MM3, MM4, MM5, and MM6) and one uveal melanoma (UM22) to produce CAR-TILs. CAR expression was detected by HER2-biotin binding at 5-16 hours post-transfection (a). MM1 TILs and CAR-TILs were co-cultured with the parental or HER2 KO 92-1 uveal melanoma cell line for 4-6 hours, followed by flow cytometry to measure degranulation (CD107a expression) (b) or IFN-γ-secreting cells (c). TILs and CAR-TILs not co-cultured with 92-1 cells (TILs alone; TA) were used as controls in b-c. Alternatively, cells were co-cultured for 48 h, and the supernatant was collected for analysis of secreted IFN-γ by ELISA (d). 92-1 cells not co-cultured with TILs (no TILs) were used as a control. Data are presented as mean ± standard deviation (SD) of duplicates. The experiments were performed twice, and representative results are shown.

Next, we co-cultured MM1 or CAR-TILs with 92-1 cells and stained them with an antibody directed against cleaved caspase-3. 92-1 cells cultured with CAR-TILs, but not TILs, were positive for intracellular cleaved caspase-3, suggesting that the cells underwent apoptosis (**Fig 2A**). We also generated patient-derived cell lines, TILs, and CAR-TILs from MM5 cells. CAR expression in autologous MM5 TILs enhanced their ability to kill melanoma cells (**Fig 2B**). Four out of five patient-derived cell lines were more sensitive to increasing amounts of CAR-TILs than to TILs in the co-culture experiments (**Fig 2C-G**). This correlated with the greater degranulation of CAR-TILs than mock-transfected TILs (**Suppl Fig S3**).

**Figure 2.**
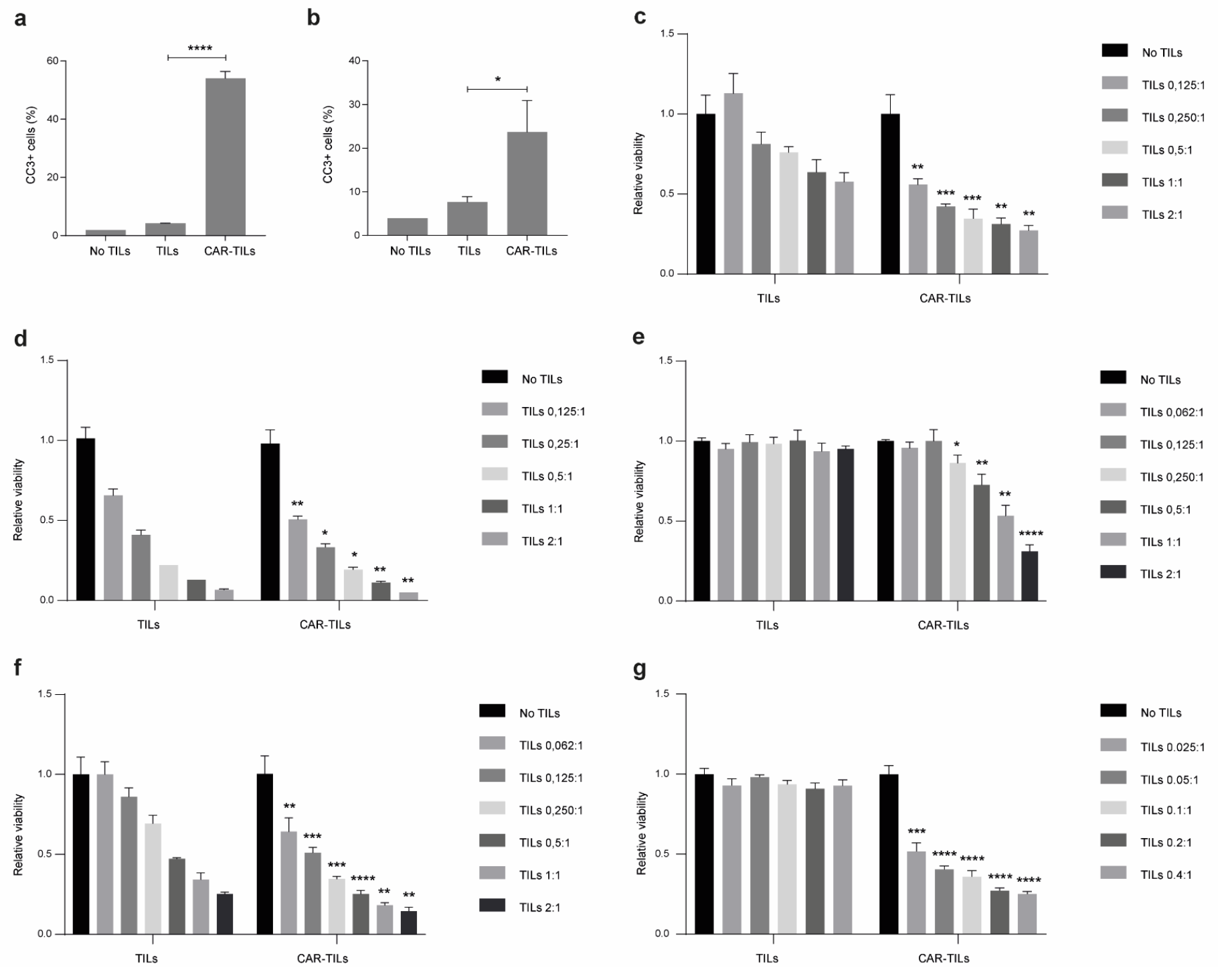
CAR-TILs kill tumor cells more efficiently than TILs. (a-b) TILs and CAR-TILs from MM1 were co-cultured with the parental or HER2 KO 92-1 uveal melanoma cell line for 24 hours, followed by cleaved caspase 3 (CC3) detection in the tumor cells using flow cytometry (a). Additionally, TILs and CAR-TILs from MM5 were co-cultured with autologous cancer cells for 24 hours, followed by cleaved CC3 detection using flow cytometry (b). Cancer cells not treated with TILs were used as a control (no TILs). (c-g) Viability of autologous cancer cells was measured by luciferase signal detected after 48 hours co-culture with increasing doses of TILs and CAR-TILs from UM22 (c), MM3 (d), MM4 (e), MM5 (f) and MM6 (g) in the indicated ratios (TILs:cancer cells). Data is presented as mean with SD of triplicates. Asterisks represent p-values of difference between similar doses of TILs and CAR-TILs. The experiments were performed twice, and representative data from one experiment is shown.

### CAR-TILs can kill melanoma cells in vivo

The mRNA transfection of CAR into TILs allowed for a fast way to assess the efficacy of CAR in TILs, but the expression was not durable. TILs kill via the binding of the T-cell receptor (TCR) to the MHC class I complex, which is loaded with peptide. The MHC complex consists of one alpha chain (encoded by *HLA A/B/C*) and one Beta-2-microglobulin (B2M) chain (encoded by *B2M*). To challenge the TILs, we deleted B2M by CRISPR/Cas9 (*B2M* KO; **Suppl Fig S4**) in a cell line from patient MM3, rendering the tumor cells resistant to killing by autologous TILs but not to anti-HER2 CAR mRNA-transfected CAR-TILs (**Fig 3A**).

**Figure 3.**
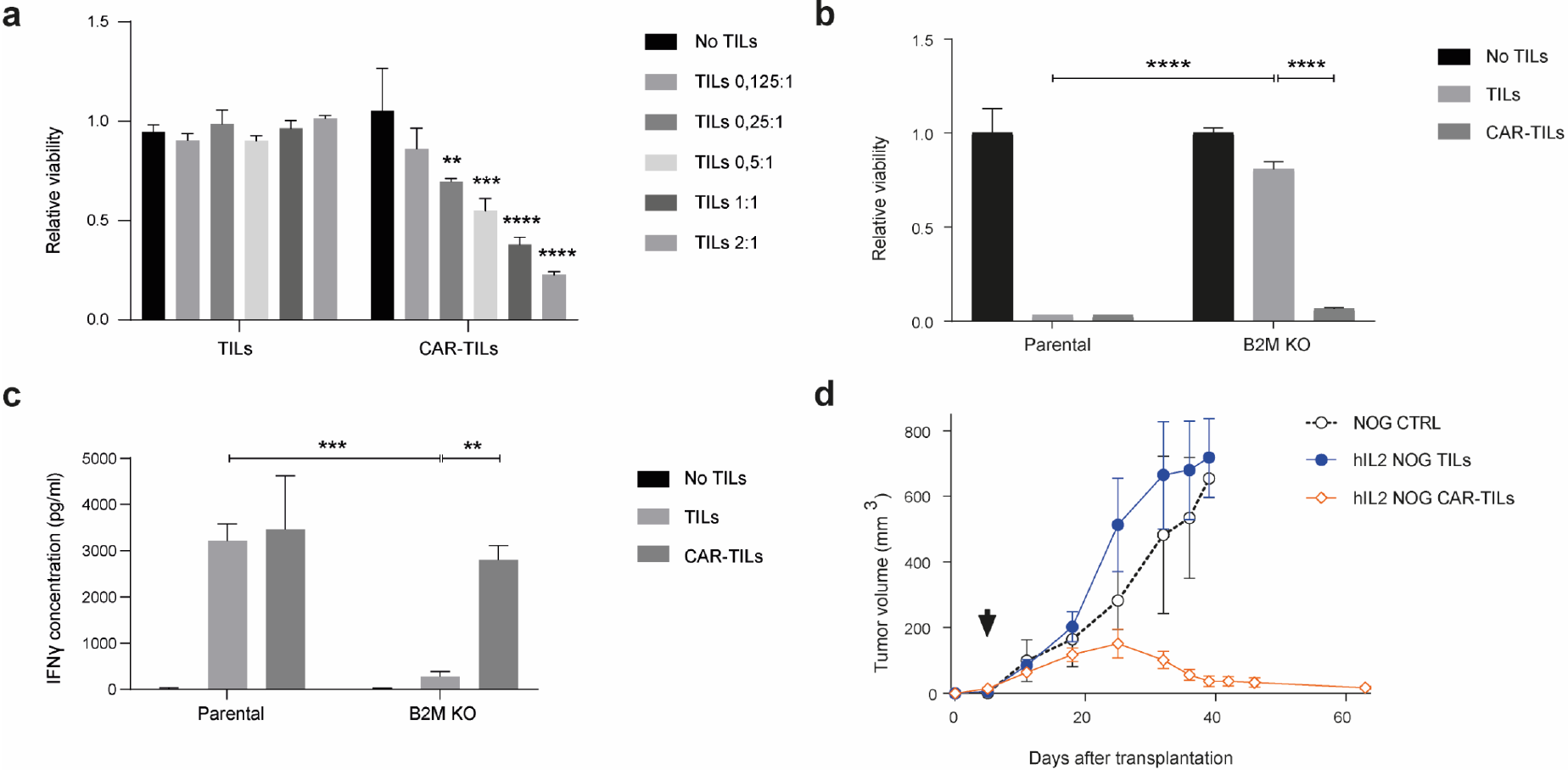
Anti-HER2 CAR-TILs eradicate autologous tumor cells refractory to TCR-mediated killing. Parental MM3 melanoma and MM3 cells with CRISPR-Cas9 disruption of β-2-Microglubulin (B2M KO, Suppl Fig 5) were cultured with autologous TILs or CAR-TILS. (a-b) Viability was measured by luciferase activity in melanoma cells after 48 h of co-culture with TILs or mRNA generated (a) or virus generated (b) CAR-TILs. (c) IFNΨ was measured in the culture supernatant from the same experiment as in b. (d) Mice bearing B2M CRISPR knockout MM3 melanoma cells were treated with PBS (n=3), 20 × 10^6^ TILs (n=3), or 20 × 10^6^ CAR-TILs (n=3). Data are presented as mean ± standard error of the mean.

To study the effect of CAR-TILs *in vivo* we used a lentivirus expressing the same anti-HER2 CAR mRNA (40). CAR expression was much lower in lentivirus-transduced cells than in those transfected with CAR mRNA and only 1-10% of melanoma cells were susceptible to lentivirus transduction (**Suppl Fig S5**). Nevertheless, this modification of MM3 TILs enabled the killing of both parental MM3 cells and *B2M* KO tumor cells *in vitro* (**Fig 3B**), which correlated with IFN-γ secretion into the medium.

We previously showed that TILs and blood-derived HER2 CAR-T cells can eradicate PDX tumor models in the human IL-2 transgenic NOG/NSG mouse strain (hIL2-NOG). We transplanted *B2M* KO MM3 cells into hIL2-NOG cells and treated them with TILs or anti-HER2 CAR-TILs. As expected from the *in vitro* experiments (**Fig 3A-C**), TILs were not able to eradicate B2M deficient melanoma in the PDXv2 model, but CAR-TILs could (**Fig 3D**). In PDX models from UM1, MM2, and MM7, the added value of lentiviral CAR expression in TILs was only visible in MM2, since unmodified TILs could eradicate tumors in UM1 and MM7 PDX models (**Suppl Fig S6**).

### <u>F</u>irst-in-<u>do</u>g (FIDO) trial suggest CAR-TIL therapy is safe and may have anti-tumoral activity in tumor bearing companion dogs

To assess the safety of anti-HER2 CAR-TILs in an immune-competent and spontaneous tumor model before the first-time-in-<u>m</u>an trial (FTIM), we conducted a first-in-<u>do</u>g (FIDO) trial that recruited four companion dogs with high-grade cancer (two with squamous carcinoma and two with melanoma). In parallel with the recruitment and treatment of dogs, we also performed several analyses of the tumors and TILs. First, we assessed the expression of HER2 in the tumors of all dogs compared to blood leukocytes, canine D17.os osteosarcoma and CF41.mg mammary cell lines. The Dog 2 tumor had the highest HER2 mRNA expression, followed by the mammary cell line and the second melanoma dog (Dog 4). The tumors of dogs 1 and 3 had similar expression as the osteosarcoma line, whereas blood leukocytes were practically negative for HER2 expression (**Fig 4A**). Immunohistochemistry demonstrated HER2 expression in all dog tumors (**Fig 4B**).

**Figure 4.**
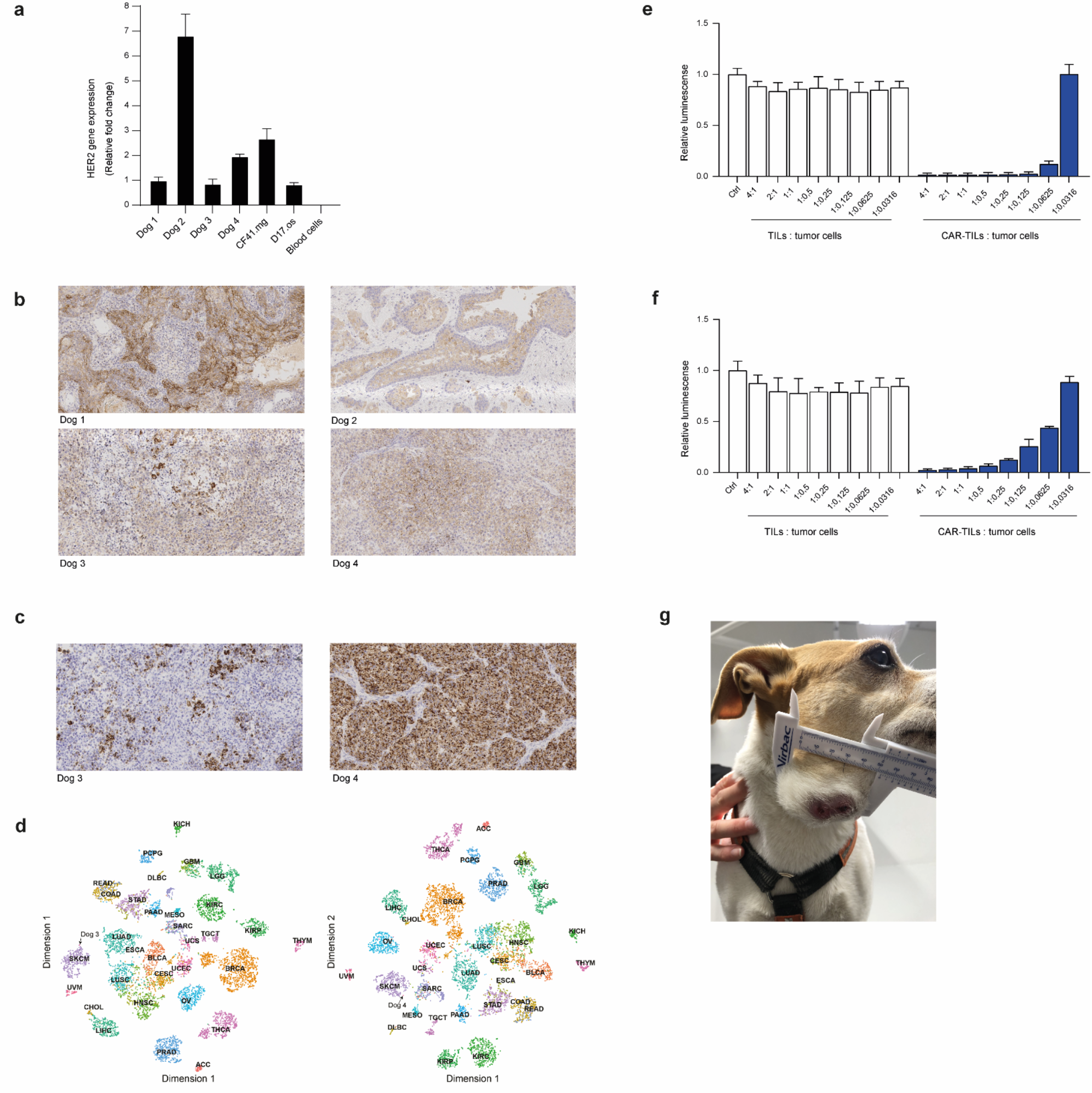
Dog tumor cells express HER2 and can be killed by anti-HER2 CAR-TILs. HER2 expression detected by qPCR (a) and immunohistochemistry (b) in tumors from four companion dogs (dogs 1-4). Canine cell lines D17.os and CF41.mg and canine PBMC were used as controls in a. (c) Expression of human melanoma marker Melan-a in tumors from dogs 3 and 4 with melanoma. (d) Classification of canine tumor biopsies from dogs 3 and 4 based on gene expression relative to TCGA cohort of > 10000 human tumor samples from 32 cancer types. The similarities between canine melanoma and samples in TCGA were visualized using tSNE dimensionality reduction. SKCM: melanoma cases. (e-f) TILs and mRNA electroporated CAR-TILs from dog 3 were co-cultured with the canine cell lines CF41.mg (e) and D17.os (f) for 24 h. The experiment was performed twice and representative data (n=3, data ± SD) from one experiment are shown. (g) Photograph of Dog 3 with metastatic oral malignant melanoma enrolled in the FIDO trial evaluating the safety of CAR-TIL treatment in companion dogs.

We recruited two dogs with melanoma, Dog 3, which had patchy expression, and Dog 4, which had uniform expression of the melanoma marker Melan-a (**Fig 4C**). To compare the transcriptomes of canine and human cancers, we used an in-house bioinformatics pipeline developed to diagnose cancer of unknown primary origin (44). Comparing gene expression of around 10 000 human tumors in The Cancer Genome Atlas (TCGA) with those of the biopsies from dogs 3 and 4 using k-nearest neighbor analysis, we found that canine melanomas were most similar to human melanoma (SKCM) of all tumors in TCGA (**Fig 4D**).

To evaluate whether our anti-HER2 CAR binder would bind to canine HER2, we transfected canine TILs with the mRNA encoding anti-HER2 CAR. We co-cultured the TILs or anti-HER2 CAR-TILs with the canine D17.os and CF41.mg cell lines labeled with firefly luciferase. Since luciferase requires ATP to glow, we were able to assess the viability of tumor cells exclusively by luciferase measurements when co-cultured with increasing amounts of TILs or CAR-TILs. None of the cell lines were sensitive to allogenic TILs, but both were killed by CAR-TILs in a dose-dependent manner (**Fig 4E-F**). The canine mammary carcinoma cell line was more sensitive than the osteosarcoma cell line, suggesting that the level of HER2 expression (**Fig 4A**) affects sensitivity to CAR-TILs.

The FIDO trial design was a dose-escalation study giving 0.1-10 million CAR-TILs/kg dog, without or with injections of human IL-2. All dogs received one dose of the lowest dose of CAR-TILs, and after safety monitoring, one or two additional doses with or without IL-2 were administered. The first two dogs had squamous cell carcinoma of the tongue and tonsils. The dogs initially tolerated the treatment well; however, the first dog developed pyometra, a common disease in female dogs of this breed. None of the other two female dogs in the study developed this disease, suggesting that it was not apparently related to treatment. The first dog experienced a slight reduction in tumor size, but this was not a durable response (**Suppl Table S1**). Since the percentage of anti-HER2 CAR positive cells was very low in the CAR-TILs of the first dog, and the main purpose of the trial was to evaluate the safety of the anti-HER2 CAR binder, we used anti-HER2 CAR mRNA transfection for the next dog. We achieved approximately 30% of CAR-positive cells. The tumor size was not accurately measurable from the back of the tongue, but there was no apparent softening or decrease in size with treatment, and the dog was euthanized upon tumor progression because of deteriorating quality of life (**Suppl Table S2**).

Dogs 3 and 4 suffer from melanoma, a debilitating disease in dogs, because it often affects eating; it rapidly develops systemic metastases and, if untreated, is lethal. We were able to generate more TILs from dogs with melanoma and were therefore able to inject both mRNA-transfected and virus-transduced CAR-TILs, as well as treatment with IL-2. Initially, the tumor grew rapidly on Dog 3 (**Fig 4G**), but a transient decrease in tumor size (PR) was observed after the third CAR-TIL injection and first IL-2 injections. The dog experienced gastrointestinal side effects, such as loss of appetite, vomiting, and diarrhea (**Table 1**), which were resolved and did not appear after subsequent reduction in the dose of reminders of IL-2 injections. After the initial decrease in tumor size, the tumor progressed, and the dog was euthanized because of tumor progression.

**Table 1.** Clinical responses to CAR-TIL treatment of Dog 3.

| Days after treatment | Treatment | Number of cells/kg | CAR expression | Tumor size (mm) | CRP (mg/l) | WBC (10<sup>9</sup>/l) | Eosinophils (10<sup>9</sup>/l) | Side effects |
| --- | --- | --- | --- | --- | --- | --- | --- | --- |
| 0 | Dose 1: CAR-TiLs | 0.1 million | 70% (mRNA) |  | <7 | 5 | 0.3 |  |
| 14 | Dose 2: CAR-TiLs | 10 million | 75% (mRNA) | 15 x 13 | <7 | 8.4 | 0.4 | Diarrhea that resolved after one day |
| 35 |  |  |  | 27 x 14 | <7 | 7.9 | 0.5 |  |
| 63 |  |  |  | 35 x 29 | <7 | 7.4 | 0.4 |  |
| 78 | Dose 3: CAR-TiLs | 10 million | 1-7% (virus) | 42 x 24 | <7 | 7.2 | 0.6 |  |
| 78 | IL-2 injections<br>1500IU q12h |  |  |  |  |  |  | Diarrhea and vomiting, decreased appetite all grade 1 |
| 84 | IL-2 injections<br>1500IU q12h |  |  | 36 x 22 | 58 | 10.5 | 1.3 | Eosinophilia grade 1 |
| 89 | IL-2 injections<br>1500IU q12h |  |  | 34 x 24 | 51 | 23.2 | 9.7 | Eosinophilia grade 1 |
| 112 |  |  |  | 52 x 44 | <7 | 11.2 |  |  |
| 133 | Euthanized due to tumor progression |  |  |  |  |  |  |  |

The fourth dog underwent surgery for subungeal melanoma with metastasis to the prescapular lymph node, and no detectable tumor was detected after surgery. Nevertheless, since local recurrence or distant metastases invariably develops within 3-6 months in these cases, we treated the dog with both the lowest and therapeutic dose of CAR-TILs in an adjuvant setting. No acute side effects were observed with CAR-TIL treatment; however, after IL-2 treatment, gastrointestinal side effects and eosinophilia resolved with treatment discontinuation and re-emerged when reinitiating treatment (**Table 2**). IL-2 treatment ceased, and one year after surgery, the dog was still tumor-free.

**Table 2.**
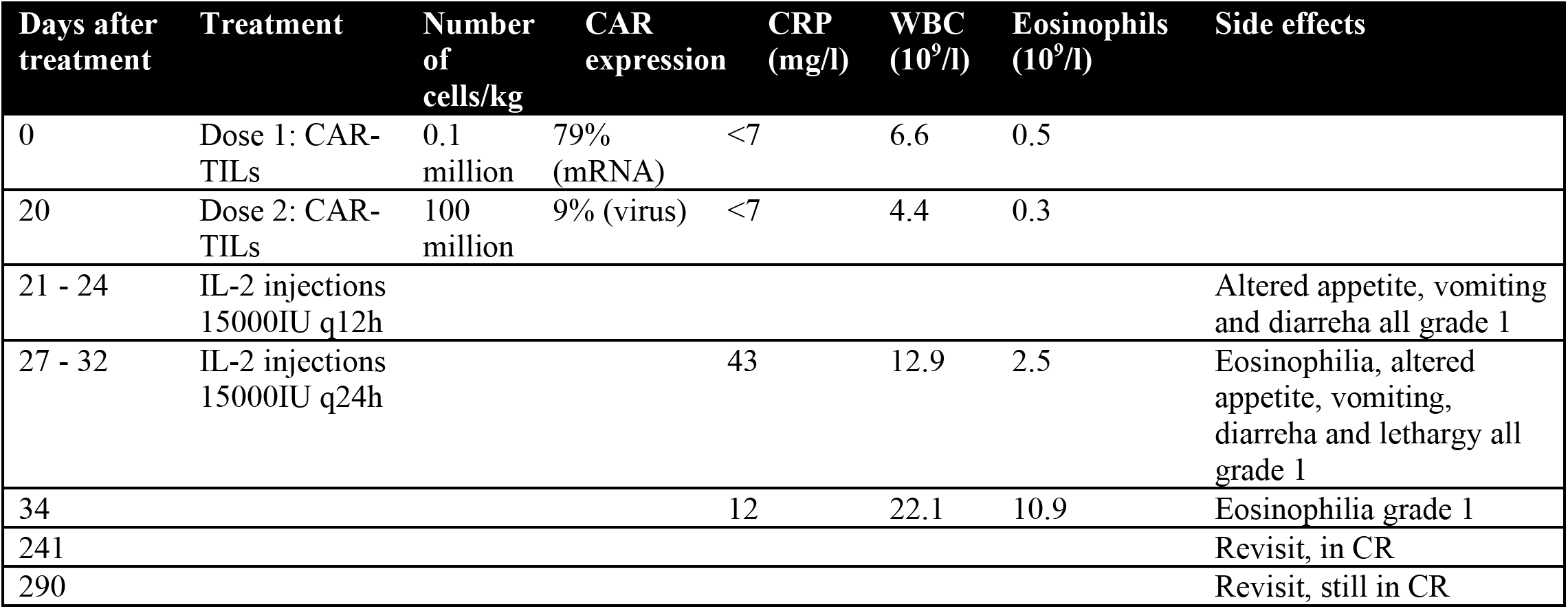
Clinical responses to CAR-TIL treatment of Dog 4.

Monitoring of safety in all four dogs throughout the treatment period did not reveal any apparent effects on blood pressure or cardiac, pulmonary, or endocrine functions. Liver and electrolyte levels were normal, but CRP was elevated in response to tumor progression, IL-2 administration, or infection (**Suppl Table S1-2** and **Table 1-2**).

## Discussion

Immunotherapies with checkpoint blockade have revolutionized the treatment of metastatic melanoma, with durable and high response rates in patients who previously had a poor prognosis. However, some tumors pose a challenge for immunotherapies because of their low antigen load or different immune evasion strategies. A great challenge in the field is how to render these tumors immunogenic.

We posed the question of whether it would be possible to enhance the effect of TIL therapy by genetic engineering with CAR. Indeed, we showed that this therapeutic approach resulted in the activation of T cells that could not kill allogenic or immune-evaded autologous tumors. We observed complete tumor regression in PDX mice from tumors that lacked B2M and were treated with CAR-TILs (but not TILs). These responses were superior to those observed in a study published while writing this manuscript (45). This can be explained by the use of different immunocompromised mouse strains. Our data confirm the importance of IL-2 supplementation for TIL and CAR-T activity *in vivo* as shown in hIL2-NOG mice(40,42). On the other hand, Mills et al. (45) reported better transduction efficiencies using CAR-expressing retroviruses, although they selected cells to ensure high expression levels. This information is valuable for advancing the concept of CAR-TILs to the clinic.

CAR-TIL treatment could be used in patients with cancers that are normally not recognized by the immune system, for instance, due to downregulation of MHC or the antigen presentation machinery, lack of suitable neoantigens, or expression of any inhibitory receptors (including but not limited to PD1 and CTLA-4). One PDX model, which is a poor responder to TIL therapy (MM2), was rendered more responsive to T-cell killing after equipping the TILs with a CAR construct. However, tumors that respond to ACT treatment with autologous TILs (e.g., UM1 and MM7 in this study) may not be eradicated more efficiently by CAR-TIL treatment. Hence, a potential target population for a trial could consist of patients with immune checkpoint inhibitor-resistant metastatic cutaneous, mucosal, or uveal melanoma.

A challenge for immunotherapy research is to utilize good preclinical models to study human tumors and to generate preclinical efficacy and safety data that can warrant the start of clinical trials. Most preclinical advances have depended on *in vitro* studies of human tumors and immune cells, mouse tumor models, and mouse immune cells. *In vitro* systems do not always recapitulate the *in vivo* setting in a satisfactory manner, highlighting the importance of using *in vivo* models to develop novel treatment strategies for cancer patients. Mouse models used to study mouse tumors and immune cells accurately recapitulate the complex truth(46), but with limitations attributed to differences between mouse and human biology. The hIL2-NOG mouse model used here (42) has the advantage of recapitulating the heterogeneous nature of TIL therapy in cancer patients, which is not possible in NOG mice. Unfortunately, for the rare form of melanoma arising in the eye, uveal melanoma, there is a lack of models, and those that we have developed (40,43) most often grow slower in NSG/NOG mice than in cutaneous melanoma.

Companion dogs are emerging models for human diseases because their etiologies are very similar (34,36). In this study, we utilized the fact that solid tumors in companion dogs express the dog variant of HER2, which was also similar to the human protein, so that we could generate CAR-expressing T cells using our human-specific construct. The primary aim of the FIDO trial was to generate safety data for a planned FTIM study, as a previous trial with HER2 CAR-T cells reported a lethal incident (47). All adverse events were mild and exclusively associated with IL-2 treatment. Toxicity was concentrated in the GI tract, with loose stool, decreased appetite, and vomiting. The eosinophilia observed has been described previously and is likely due to the release of IL-5 and GM-CSF release by IL-2 stimulated CD4 positive lymphocytes (48). Reduced dosing of IL-2 resulted in control of toxicity, which was completely resolved after termination of IL-2 administration.

In the safety FIDO trial, we also observed some signs of anti-tumoral activity, as evidenced by a near partial response in a dog with melanoma and potentially a delay in recurrence in a second dog. We are cautiously optimistic about these early signs of efficacy, although mindful of the pitfalls and caveats of the experiments. First, since our mouse data comparing the efficacy of TILs and CAR-T cells in NOG vs. hIL2-NOG mice undeniably indicated that IL-2 is essential for the efficacy of these therapies, we treated the dogs with subcutaneous injections of IL-2. It is therefore not inconceivable that some of the therapeutic effects observed were mediated by this factor, or that the gastrointestinal adverse events resulted in metabolic effects on the tumor. Second, we could not detect CAR-TILs in the blood of dogs using PCR. This could be due to the CAR-TILs not surviving or expanding, but it could also be due to them leaving the blood stream because they express homing receptors for inflamed areas, such as tumors (41,42). A future study protocol will need to include metastasectomy or biopsy after CAR-TIL therapy to evaluate whether CAR-TILs reach the target, as they commonly do in humans.

In conclusion, we present a novel approach to treat metastatic cancer in mice and dogs by combining CAR-T cell therapy and adoptive T cell transfer, called CAR-TIL treatment. CAR-TILs elicited complete eradication of tumors in immune-humanized PDX mouse models of melanoma, even in the absence of antigen presentation, indicating the potential usefulness of this strategy for treating melanomas that do not respond to existing therapies. This regimen should be tested in patients with metastatic melanoma to assess the potential of this therapy. Since owners and veterinarians of companion dogs with melanoma have difficulty accessing effective immunotherapies when they progress on approved therapies, the data here provide an interesting avenue for future trial activities. Companion dogs also contribute to 3R by reducing (and long-term replacing) the use of healthy beagles for safety studies and refining, since companion dogs have spontaneous tumor formation and therefore are more representative of patients in FTIM studies than using beagles.

## Methods

### Human patient samples

Tumor samples were obtained from patients treated at the Department of Surgery, Sahlgrenska University hospital, Gothenburg, Sweden following informed consent (Regional Human Ethics Board of Västra Götaland, Sweden approval #288-12 and #44-18). Tumor cells were extracted from tumor samples and used for patient-derived xenograft establishment, as previously described (49). Young TILs (y-TILs) were extracted from the same tumor samples, cultured, and expanded, as previously described (42).

### Cell experiments

Patient-derived melanoma cell lines were maintained in RPMI with 10% fetal bovine serum and were either described previously [UM22, MM2, MM3, MM4 (42,43)] or generated by culturing melanoma samples in RPMI with 10% FCS (MM5, MM6). 92-1 Uveal melanoma cells were a kind gift from the European Searchable Tumor Line Database and Cell Bank, and the HER2 knockout line was established previously (40). The canine tumor cell lines D17.os and CF41.mg were purchased from ATCC.

For viability experiments, melanoma cells were infected with a lentivirus made using pHAGE-PGK-GFP-IRES-LUC-W (Addgene # 46793) containing coding sequences for green-fluorescent protein and firefly luciferase. To assess TIL killing, patient-derived melanoma cell lines expressing luciferase were plated at 20.000 cells/well in black 96-well plates (Corning) and cultured in the presence or absence of different ratios of TILs or CAR-TILs per well. After 48 h, the medium was aspirated for IFN-γ secretion analysis using an ELISA kit (Diaclone), and the viability of the cancer cells was assessed by measuring luminescence with a GloMax Discover plate reader (Promega) or an IVIS Lumina III XR (Perkin-Elmer) after adding luciferin (150 μg/ml) to the cells. For degranulation analysis, TILs and CAR-TILs were co-cultured with cancer cells for 4-6 hours in RPMI supplemented with human AB serum (Sigma-Aldrich) and CD107a antibody (clone H4A3, BD Pharmingen), followed by washing in PBS and detection of bound CD107a antibody by flow cytometry (BD Accuri C6 Plus). For the IFN-γ secretion assay, the degranulation assay was followed by additional staining procedures according to the manufacturer’s instructions (Miltenyi Biotech), and bound IFN-γ was detected using flow cytometry. For detection of cleaved caspase 3 (CC3), cancer cells were co-cultured with TILs and CAR-TILs for 24 h, followed by fixation and permeabilization using Cytofix/Cytoperm solution for 25 min (554714, BD Biosciences), washed twice with supplemented Wash/Perm buffer (554714, BD Biosciences), and stained with a CC3 antibody (clone C92-605, BD Pharmingen) at 1:10 dilution in Wash/Perm buffer. Finally, the cells were washed and analyzed using a BD Accuri C6 Plus flow cytometer equipped with BD Accuri C6 software.

For CRISPR/Cas9 inactivation of *B2M*, the Cas9:crRNA:tracrRNA ribonucleoprotein (RNP) complex was assembled according to the manufacturer’s recommendations (IDT DNA) and transfected into the cells using Lipofectamine RNAiMAX reagent (Invitrogen). Negative cells were sorted based on the absence of B2M-PE antibody staining (clone 2M2, BioLegend) using magnetic separation with PE-beads (Miltenyi), confirmed negative by staining with the same antibody, and analyzed using an Accuri C6 flow cytometer (BD) equipped with the BD Accuri C6 software.

### Human TIL and CAR-TIL expansion

For rapid expansion (REP) of TILs, yTILs (1×10^5^) were mixed with irradiated (40 Gy) feeder cells (20×10^6^), CD3 antibody (clone OKT3, 30 ng/ml, Miltenyi), and a medium containing 50% RPMI and 50% AIM-V (Invitrogen) supplemented with 10% human AB serum (Sigma-Aldrich) and 6000 IU/ml IL-2 (Peprotech). CAR-TILs were produced by transducing the TILs on days 1 and 2 of rapid expansion (REP) with a lentiviral vector encoding the HER2 CAR construct in the presence of Vectofusin-1 (Miltenyi). After five days of culture at 37 °C in 5% CO_2_, half of the medium was replenished. From day 6 onwards, the flasks were inspected daily and split when necessary to maintain cell densities of approximately 1-2×10^6^/ml. After 14 days in culture, the cells were harvested, resuspended in PBS with 300 IU/ml of IL-2, and intravenously transplanted into mice (20×10^6^ cells per mouse in 100 μL).

Alternatively, REP-TILs were transfected with anti-HER2 CAR mRNA via electroporation using a 4D nucleofector (Lonza). This mRNA was generated by PCR amplification of the coding sequence of anti-HER2-CAR (40) using primers that carried a T7 RNA polymerase recognition sequence at the 5’-end. The resulting PCR product was used in an *in vitro* transcription reaction with the T7 mScript Standard mRNA Production System (Cellscript). The TILs were resuspended in P3 primary cell solution with supplement (Lonza), and mRNA was added before pulsing using the DN100 program.

### CAR detection

For qPCR detection, genomic DNA was prepared from TILs and CAR-TILs 10-14 days after the start of REP by lysing in Direct PCR Lysis Reagent (Nordic BioSite) and proteinase K. Quantitative PCR was performed in triplicate using a qPCR SyGreen mix (Techtum Lab AB), and the PCR reaction was performed with a CFX cycler (Bio-Rad). Data analysis comparing ΔCT values normalized to a reference gene (β-actin) was performed to determine the CAR copies/cell.

For analysis of HER2 protein binding capacity of CAR-TILs, 100.000 cells were incubated with 1 μg biotinylated HER2 protein (Abcam) for 30 min at 4 °C, followed by incubation with an allophycocyanin-conjugated streptavidin antibody (Jackson Immuno Research) for 25 min at 4 °C, and flow cytometry analysis was performed using an Accuri C6 flow cytometer (BD) equipped with the BD Accuri C6 software.

### Mouse experiments

All mouse experiments were performed in accordance with EU Directive 2010/63 (Regional Animal Ethics Committee of Gothenburg #2014-36, #2016-100 and #2018-1183). Non-obese diabetic-severe combined immune-deficient interleukin-2 chain receptor γ chain knockout mice (NOG mice, Taconic, Ry, Denmark) and human IL-2 transgenic NOG (*hIL2*-NOG) mice (Taconic) were used for the engraftment of tumor samples. Tumor size was monitored by caliper measurements, alternatively bioluminescent signals, using the IVIS imaging system (Perkin-Elmer). When tumor growth was confirmed by two consecutive measurements, *hIL2*-NOG mice were treated with autologous human tumor-infiltrating lymphocytes (TILs; 20 million per mouse) by intravenous injection into the tail vein. NOG mice served as untreated controls.

### Canine CAR-TIL expansion

For canine yTIL expansion, 2-3 mm3 tumor pieces were cut and placed in a 24-well plate containing RPMI medium supplemented with 10% human AB serum (Sigma-Aldrich), 6000 IU/ml human recombinant IL2 (Peprotech), 1 mM sodium pyruvate (Gibco), and 50 uM 2-Mercaptoethanol (Gibco). After 7 days, the yTILs were harvested, washed in PBS, and resuspended in PBS supplemented with 10% fetal bovine serum. Cells were stained with a CD5-PE antibody (clone YKIX322.3, eBioscience) diluted 1:50 for 20 min at 4 °C, washed in PBS, and stained with PE microbeads (Miltenyi) according to the manufacturer’s protocol. Finally, cells were washed and CD5 positive cells were sorted using MACS separation with LD columns (Miltenyi) using a QuadroMACS Separator (Miltenyi) according to the manufacturer’s instructions. CD5 positivity was confirmed by analyzing PE expression using flow cytometry (BD Accuri C6 Plus). CD5 positive yTILs were either cryopreserved for later expansion or directly expanded using a rapid-expansion protocol (REP). For REP of canine TILs, CD5 positive yTILs (1×10^5^) were mixed with 20 × 106 irradiated (40 Gy) feeder cells from healthy dog blood donors (three different donors) and human CD3 antibody (clone OKT3, 30 ng/ml, Miltenyi) and expanded for 14 days in RPMI supplemented with 10% human AB serum (Sigma-Aldrich), 6000 IU/ml human IL2 (Peprotech), 1 mM sodium pyruvate (Gibco), and 50 uM 2-Mercaptoethanol (Gibco). For CAR-TIL production, cells were electroporated with anti-HER2 CAR mRNA after expansion and used to treat the patient the next day. Alternatively, anti-HER2 CAR lentivirus was added after 1, 2, and 12 d of culture.

### First-in-dog (FIDO) trial

Four client-owned dogs with HER-2 positive aggressive tumors were recruited at the University Animal Hospital (Uppsala, Sweden) between November 2019 and April 2021. All the dogs had recurrent spontaneous cancers with metastases. The CAR-TIL FIDO study and sample collection were approved by the Swedish Animal Ethical Committee and Swedish Animal Welfare Agency (#2019-2435). Written consent was obtained from dog owners.

The study design was as follows: At the initial visit, blood samples were collected and hematology and biochemistry were analyzed. The dogs were screened with a whole-body CT scan (dogs 1 and 4) or thoracic X-rays and ultrasound of the local lymph nodes and abdomen (dogs 2 and 3). As soon as possible, surgical extirpation of the primary tumor was performed in three dogs (dog 1-3) and of the metastasized local lymph nodes in one dog (dog 4). The tumors were sent to a pathology laboratory for diagnostic purposes. Parts of the tumors were sent in sterile PBS buffer with Primocin^®^ 1:500 to the laboratory in Gothenburg for expansion of autologous TILs. The TILs were modified, grown, and harvested as described below.

Before the first dose of CAR-TILs, each dog was treated for 10-14 days with oral Toceranib phosphate (Palladia^®^) 2.3-2.5 mg/kg *qd* to suppress regulatory T-cells. Treatment was stopped 5 days before CAR-TIL transfusion. The dogs were sedated with subcutaneous injections of medetomidine:butorphanol (0.01 mg/kg:0.1 mg/kg). Transfusion of CAR-TILs was performed via a peripheral venous catheter for 30 min using a protocol that included monitoring vital signs (body temperature, breathing, mucous membrane heart rate, blood pressure, and pulses). After transfusion, the sedated dogs were reverted with an intramuscular injection of atipamezol (0.05 mg/kg). Dogs stayed at the clinic for at least one hour after transfusion for supervision. Three dogs were treated with amoxicillin 9.5-11.5 mg/kg bidaily (BID) for ten days and one (dog 2) was treated with clindamycin 12 mg/kg BID for ten days, after each CAR-TIL treatment.

Blood samples were collected at visits for treatment and revisits. A complete blood count, standard clinical chemistry profile, and immunoglobulin gel electrophoresis were performed. Response to therapy was categorized in accordance with veterinary-adjusted RECIST criteria (50) as CR (complete regression of measurable soft tissue disease), PR (partial response of at least 30% reduction in the sum of diameters of target lesions, taking as reference the baseline sum), PD (progression of disease by either the appearance of one or more new lesions or at least a 20% increase in the sum of diameters of target lesions, taking as reference the smallest sum on study), and SD (less than 30% reduction or 20% increase in the sum of diameters of target lesions, taking as reference the smallest sum of diameters while on study). The best overall response was defined as the best response recorded from the start of treatment to disease progression or recurrence. Macroscopic tumor lesions were measured using a caliper and documented using photographs. In addition, dogs were followed up with diagnostic imaging (computed tomography, radiography, and/or ultrasound), using the most suitable modality for each case. Adverse events were graded according to VCOG-CTCAE version 2(51).

### Protein analysis

For immunohistochemistry, dog tumor tissues were fixed in 4% formalin, dehydrated, and embedded in paraffin. Sections (4 μm were mounted onto positively charged glass slides and dried at 60° C for 1 h. The slides were rehydrated, and antigen retrieval was performed by heat-induced epitope retrieval (HIER) in Dako PT Link with a high pH buffer (Dako). The staining was performed with an autostainer (Autostainer Link 48, Dako) using the following protocol: Endogenous Enzyme Block for 5 min (FLEX Peroxidase Block, Dako), primary antibody (HER2 A0485 and Melan-A IR633; both from Dako) staining for 60 min at room temperature, secondary reagent staining (FLEX + Rabbit LINKER, K8009, Dako) for 15 min, FLEX HRP for 20 min, diaminobenzidine (DAB) chromogen development for 10 min, and counterstaining with hematoxylin for 12 min. The slides were dehydrated and mounted using a Pertex.

### Statistical analyses of experimental data

Statistical analysis was performed using GraphPad software, ordinary one-way ANOVA, multiple comparisons with Tukey’s correction (fig 1 and fig 3 b-c), and alternatively unpaired T tests (fig 2, and 3 a). P values are represented as * P < 0.05; ** P < 0.01; *** P < 0.001; and **** P < 0.0001. All mouse experiments contained 3-4 mice per group.

### Single cell gene expression analysis

Single-cell RNA-seq from two recent studies (24,43) was used to compare the gene expression profiles of CD8^+^ and CD4^+^ T cells found in tumors (TILs) and blood from patients with uveal melanoma. Alignment and estimation of gene expression levels were performed using Cell Ranger (v. 3.0.2, 10x Genomics). The specific commands used were *cellranger count* (with the 10x Genomics version of the GRCh38 reference transcriptome; v. 3.0.0) and *cellranger vdj* (with the 10x Genomics GRCh38 VDJ reference dataset; v. 2.0.0). After identifying cell types, the remaining analyses were performed using the *Seurat* R package (v. 4.0.3) (52), and data were imported and normalized using the *NormalizeData* function with default settings. Cells predicted to be duplicates were excluded from statistical tests using R package *DoubletFinder* (v. 2.0.3, parameters: PCs=1:15, pN=0.25). Cells with more than one TCR alpha or beta chain were also excluded; however, not all cells were duplicates. An approach described by Karlsson et al. (43) was used to classify the cell types in both datasets. Differential expression was assessed using the Seurat FindMarkers function (test.use = “LR,” logfc.threshold=-Inf) between the same cell type in the two datasets. Note that the batch effects cannot be excluded from the analysis. FindMarkers were run at three different times with or without accounting for the cell cycle (derived from the Seurat function CellCycleScoring) and sex, and only genes commonly identified as differentially expressed in all three analyses were retained (**Suppl Table 3**).

### RNA-sequencing

RNA was prepared from companion dog tumors using the RNA/DNA kit from Qiagen. RNA was sequenced at the Clinical Genetics Center at Sahlgrenska University Hospital using a Novaseq sequencer. Reads were aligned to the CanFam3.1 genome (http://ngi-igenomes.s3.amazonaws.com/igenomes/Canis_familiaris/Ensembl/CanFam3.1/Sequence/WholeGenomeFasta/genome.fa) with STAR (v. 2.7.10a)(53), with a matching reference genome annotation supplied from AWS iGenomes (http://ngi-igenomes.s3.amazonaws.com/igenomes/Canis_familiaris/Ensembl/CanFam3.1/Annotation/Genes/genes.gtf), using the parameters “--twopassMode Basic” and setting “—sjdbOverhang” equal to read length – 1. Gene expression levels were quantified from name-sorted and non-duplicate marked alignment files using htseq-count (v. 1.99.2)(54), with the parameters “-s reverse -m intersection-strict”.

### Transcriptomic classification

TCGA data, downloaded and processed as described previously (44), were used for the classification of canine tumors relative to human cancer types. Pairwise Spearman correlation coefficients were calculated between our sample and each TCGA sample for all coding genes. Classification was performed using a k-nearest neighbor approach based on these correlation coefficients, using *k* = 6, as previously found to be optimal based on leave-one-out cross-validation in TCGA cohort (44).

## Acknowledgement

We wish to thank Carina Karlsson for technical support and Larissa Rizzo, Sofia Stenqvist, and Mona Svedman for assistance with animal experiments. We want to thank NIH/NCI preclinical repository for the kind donation of human IL-2.

This work was supported by the Swedish Cancer Society, Swedish Research Council, EU Horizon 2020 (ERA PerMed), Region Västra Götaland (ALF-grant), Knut and Alice Wallenberg Foundation, Sjöberg Foundation, Familjen Erling Persson Foundation, IngaBritt and Arne Lundberg Foundation (to JAN), Assar Gabrielsson Foundation, and Sahlgrenska Universitetssjukhusets stiftelse (Sahlgrenska University Hospital, Gothenburg) (to EF). SS and HR were supported by the Johansson Family Swedish Boxer Club Cancer Donation, and AH was supported by the Jane and Aatos Erkko Foundation. JAN, LMN, and JK were supported by the Kirkbride Melanoma Discovery Lab at the Harry Perkins Institute of Medical Research.

## Data Access Statement

All sequencing data has been deposited to European Genome Archives and made available under controlled data access.

## Tables and Figures

Table 1 Clinical responses to CAR-TIL treatment of Dog 3 (Billy)

| Days | Treatment | Cells<br>#/kg | CAR<br>expression | Tumor<br>size<br>(mm) | CRP<br>(mg/l) | WBC<br>(x10<sup>9</sup>/l) | Eosinophils<br>(10<sup>9</sup>/l) | Side effects |
| --- | --- | --- | --- | --- | --- | --- | --- | --- |
| 0 | Dose 1:<br>CAR-TILs | 1<br>million | 70%<br>(mRNA) |  | <7 | 5 | 0,3 |  |
| 14 | Dose 2:<br>CAR-TILs | 100<br>million | 75%<br>(mRNA) | 15 x 13 | <7 | 8,4 | 0,4 | Diarrhea<br>resolved<br>after one<br>day |
| 35 |  |  |  | 27 x 14 | <7 | 7,9 | 0,5 |  |
| 63 |  |  |  | 35 x 29 | <7 | 7,4 | 0,4 |  |
| 78 | Dose 3:<br>CAR-TILs | 100<br>million | 1-7%<br>(virus) | 42 x 24 | <7 | 7,2 | 0,6 |  |
| 78 | IL-2<br>injections<br>1500IU<br>q12h |  |  |  |  |  |  | Diarrhea,<br>vomiting,<br>decreased<br>appetite, all<br>grade 1 |
| 84 | IL-2<br>injections<br>1500IU<br>q12h |  |  | 36 x 22 | 58 | 10,5 | 1,3 | Eosinophilia<br>grade 1 |
| 89 | IL-2<br>injections<br>1500IU<br>q12h |  |  | 34 x 24 | 51 | 23,2 | 9,7 | Eosinophilia<br>grade 1 |
| 112 |  |  |  | 52 x 44 | <7 | 11,2 |  |  |
| 133 | Euthanized<br>due to<br>tumor<br>progression |  |  |  |  |  |  |  |

Tumor size, C reactive protein (CRP), white blood cell count (WBC), eosinophils and side effects were monitored.

Table 2 Clinical responses to CAR-TIL treatment of Dog 4 Nova.

| Days | Treatment | Cells<br>#/kg | CAR<br>expression | CRP<br>(mg/l) | WBC<br>(x10<sup>9</sup>/l) | Eosinophils<br>(10<sup>9</sup>/l) | Side effects |
| --- | --- | --- | --- | --- | --- | --- | --- |
| 0 | Dose 1:<br>CAR-TILs | 1<br>million | 79%<br>(mRNA) | <7 | 6,6 | 0,5 |  |
| 20 | Dose 2:<br>CAR-TILs | 100<br>million | 9% (virus) | <7 | 4,4 | 0,3 |  |
| 21 - 24 | IL-2<br>injections<br>15000IU<br>q12h |  |  |  |  |  | Altered appetite,<br>vomiting and<br>diarrhea, all<br>grade 1 |
| 27 - 32 | IL-2<br>injections<br>15000IU<br>q24h |  |  | 43 | 12,9 | 2,5 | Eosinophilia,<br>altered appetite,<br>vomiting,<br>diarrhea and<br>lethargy, all<br>grade 1 |
| 34 |  |  |  | 12 | 22,1 | 10,9 | Eosinophilia<br>grade 1 |
| 241 |  |  |  |  |  |  | Revisit, in CR |
| 290 |  |  |  |  |  |  | Revisit, still in<br>CR |

**Supplemental Figure S1.**
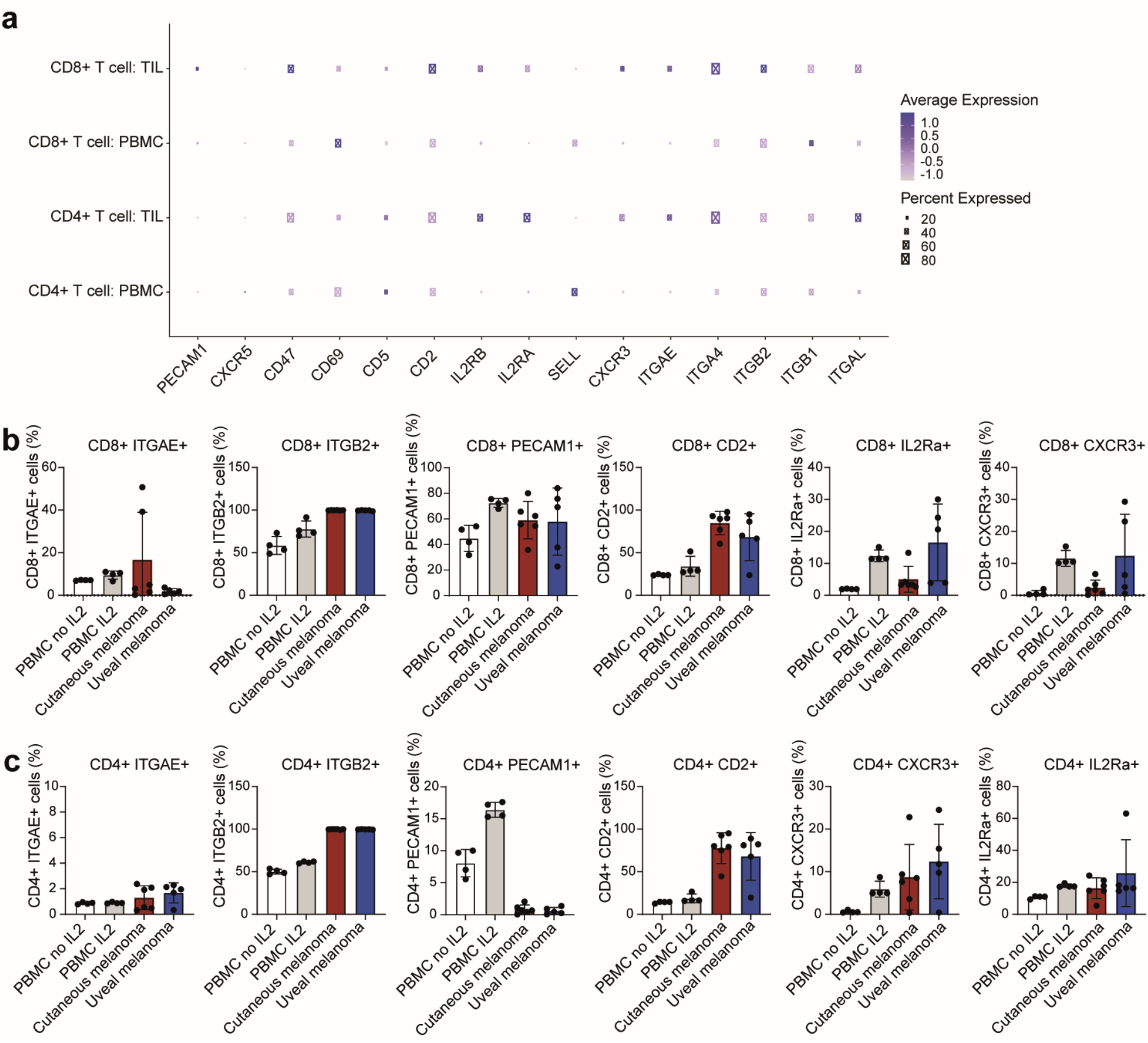
Differential expression of chemokine and adhesion between TILs and blood-derived T cells. (a) Differentially expressed genes at the single-cell level, in the categories of chemokine receptors, ligands or adhesion molecules, between CD8^+^ and CD4^+^ TILs and blood T cells (PBMC) obtained from uveal melanoma patient samples. Tests were carried out with the Seurat FindMarkers function, using logistic regression, adjusting for sex and cell cycle scores as described in Methods. (b-c) 6 of the differentially expressed genes were also analyzed with flow cytometry in both CD4+ (b) and CD8+ cells (c) coming from PBMC, skin melanoma or uveal melanoma.

**Supplemental Figure S2:**
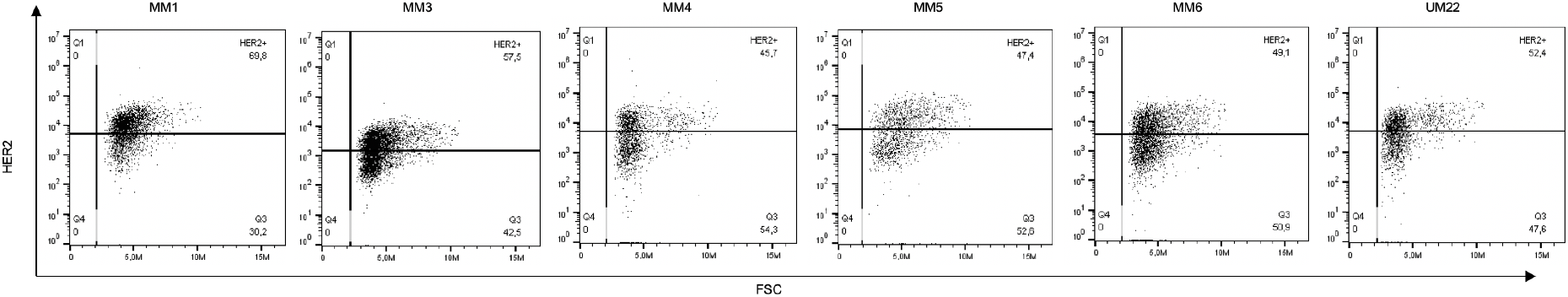
HER2 CAR-expression after electroporation. Flow cytometry data showing HER2 CAR expression after HER2 CAR mRNA electroporation by detecting bound biotinylated HER2 protein. Representative plots for samples MM1, MM3, MM4, MM5, MM6 and UM22.

**Supplemental Figure S3.**
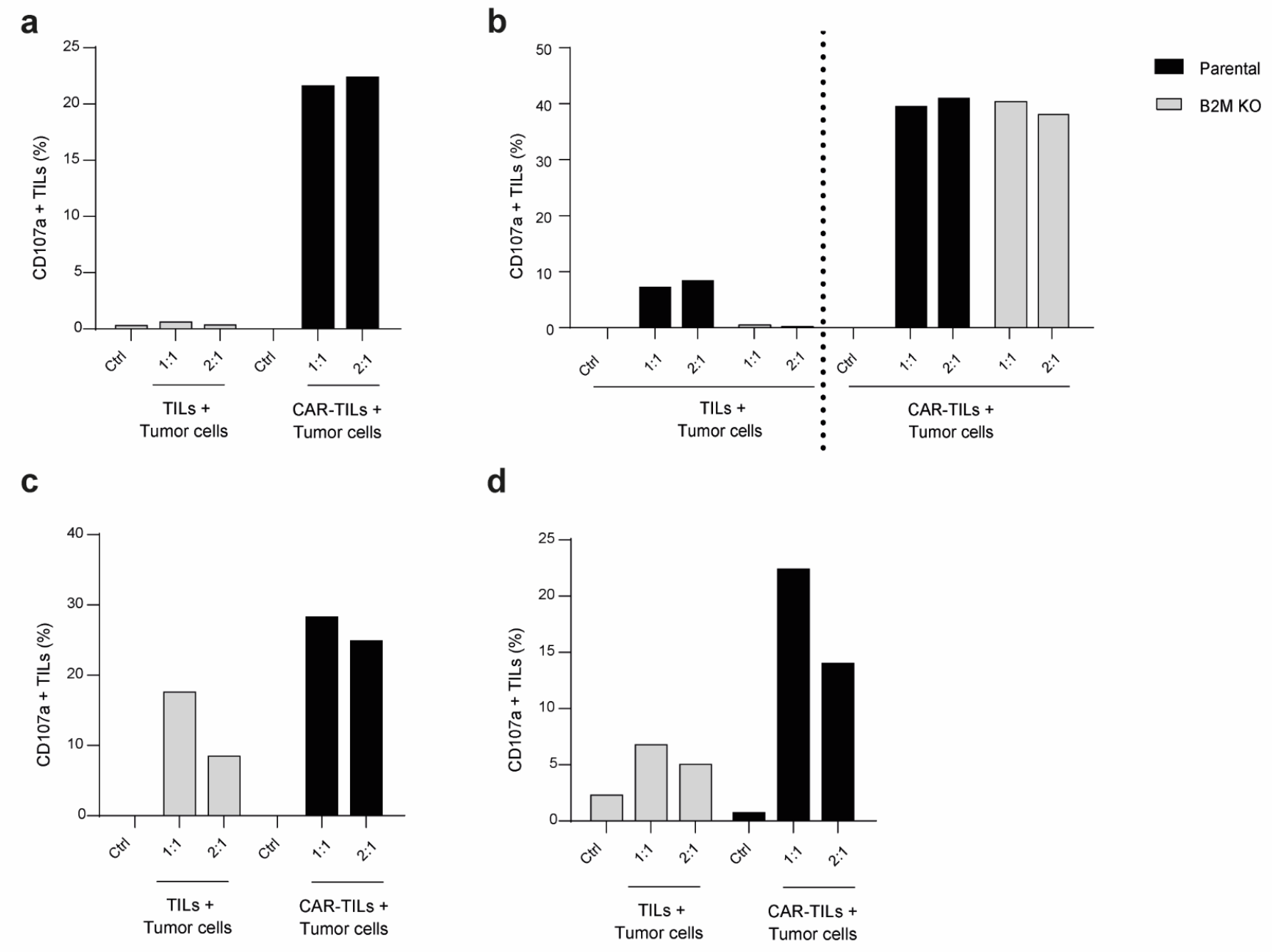
HER2 CAR-TILs degranulate more potently than TILs on autologous tumor cells. Degranulation detected by CD107a staining in TILs and CAR-TILs after 4-6 hours co-culture with autologous tumor cell lines in UM22 (a), MM3 (b), MM4 (c) and MM5 (d). The analysis was performed in singlets and twice, representative data is shown from one of the experiments.

**Supplemental Figure S4.**
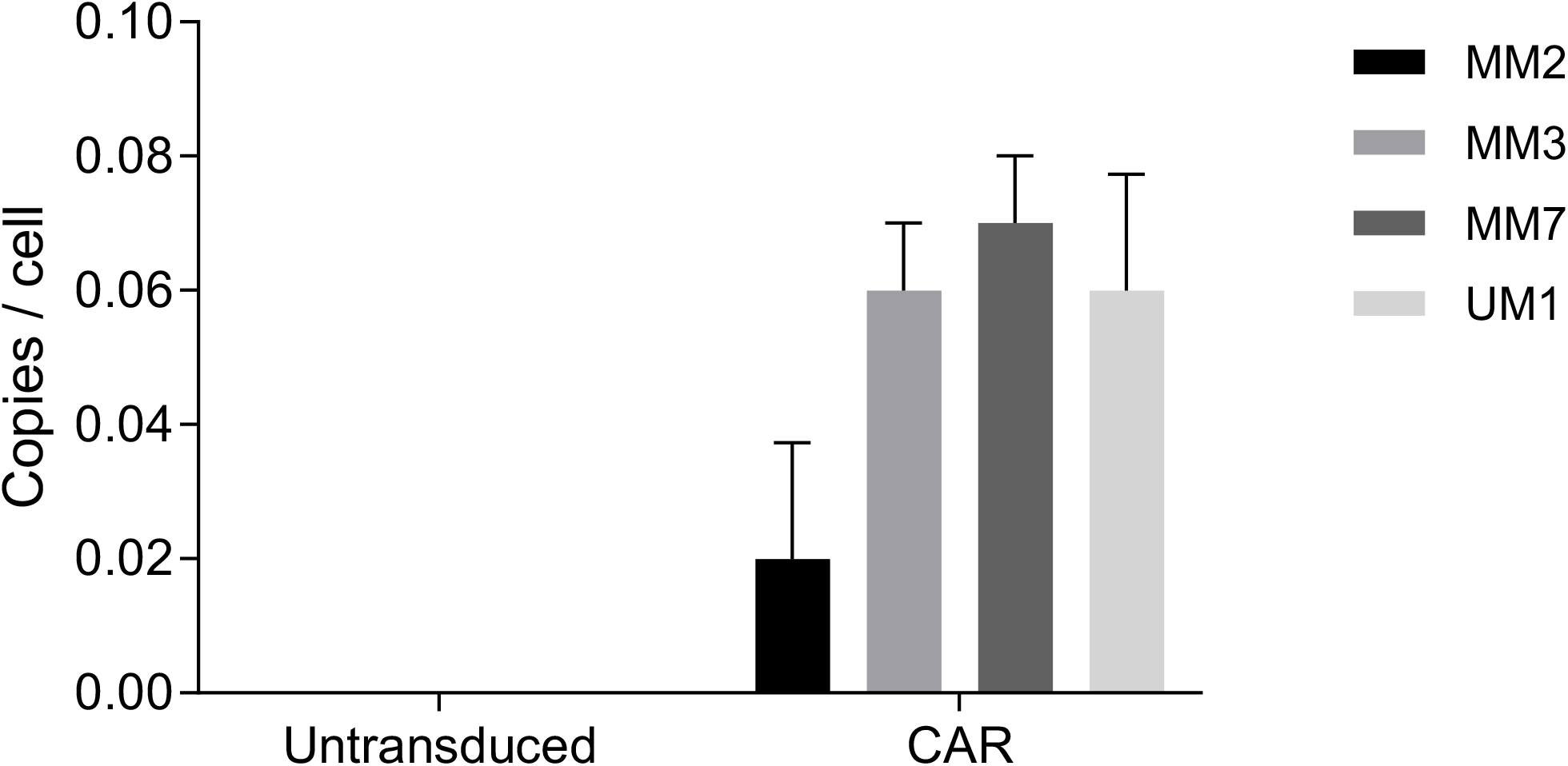
Expression of HER2 CAR in melanoma TILs. CAR-expression in TILs (untransduced) and CAR-TILs (CAR) detected by qPCR in samples MM2, MM3, MM7 and UM1. The data is presented as mean with SD of 3 replicates.

**Supplemental Figure S5.**
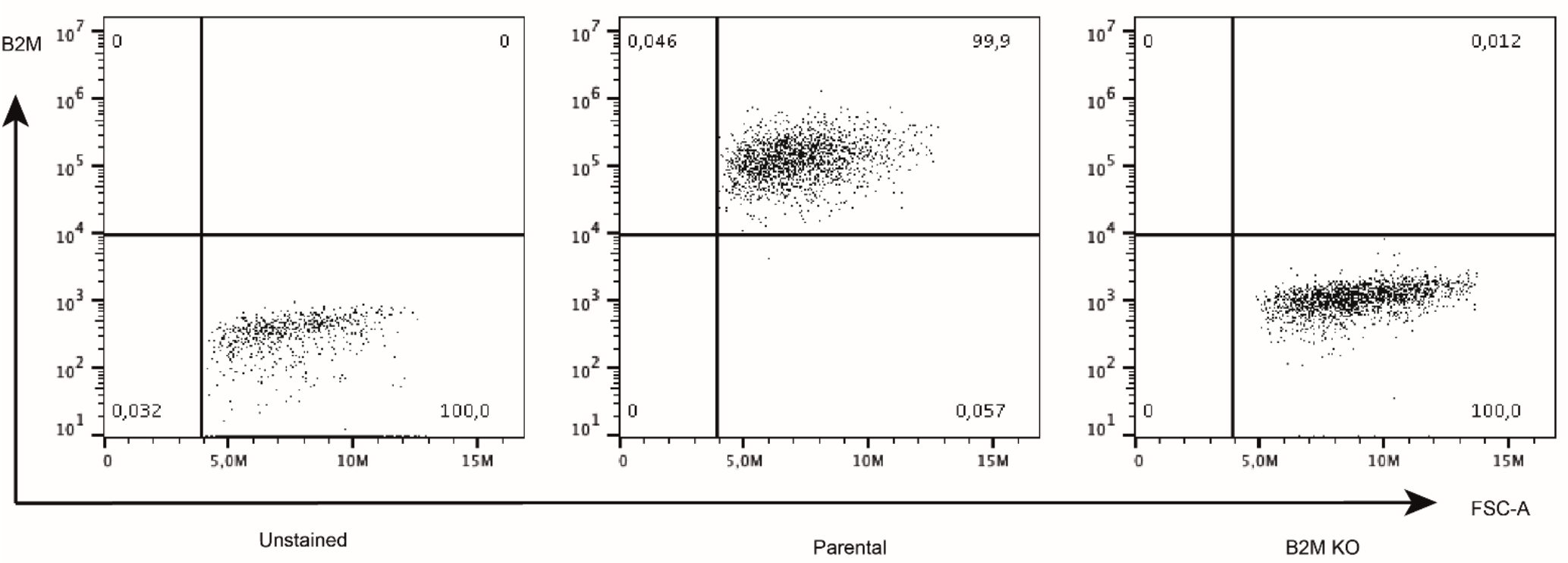
B2M KO analysis. CRISPR/Cas9 technology was used for genetic disruption of *B2M*. Cells were stained with a B2M-PE antibody and separated with magnetic sorting based on negative binding to PE-beads, and subsequently further selected by killing off B2M positive cells *in vitro* for 48 hours with autologous TILs. The resulting B2M negative cell population (*B2M* KO) and parental cells (wt) were stained with a B2M antibody and analyzed using flow cytometry.

**Supplemental Figure S6.**
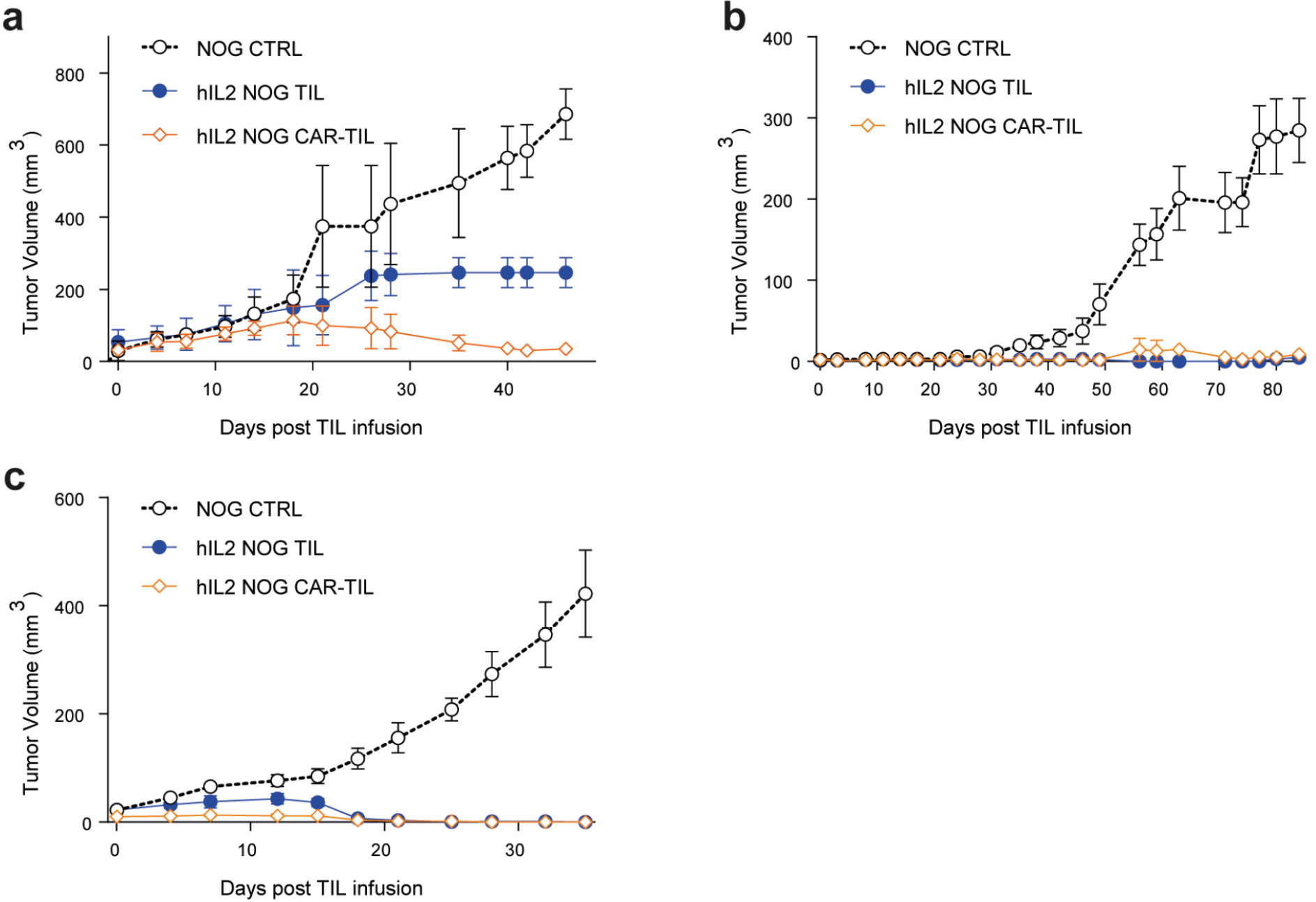
CAR-TILs can be used to treat autologous melanoma xenografts in IL2 transgenic NOG mice. TILs and lentiviral transduced CAR-TILs were produced from 3 melanoma samples and used to treat tumor-bearing mice. Two samples were from cutaneous melanoma MM2 (a) and MM7 (c) and one was from uveal melanoma UM1 (b).

**Supplemental Table 1.** Clinical responses to CAR-TIL treatment of Dog 1 (Juni)

| Days from treatment | Treatment | Number of cells/kg | CAR expression | Tumor size (mm) | CRP (mg/l) | WBC (10<sup>9</sup>/l) | Side effects |
| --- | --- | --- | --- | --- | --- | --- | --- |
| 0 | Dose 1: CAR-TILs | 0,1 million | 0,5-1,6% (virus) | 60 x 50 | <7 | 5,2 |  |
| 8 |  |  |  | 45 x 48 | <7 | 5,8 |  |
| 28 | Dose 2: CAR-TILs | 1 million | 0,4% (virus) | 40 x 40 | <7 | 10,5 |  |
| 50 |  |  |  |  | 265 | 2,7 |  |
| 51 | Euthanized due to complications to an acute perforated pyometra |  |  |  |  |  | (none reported) |

**Supplemental Table 2.**
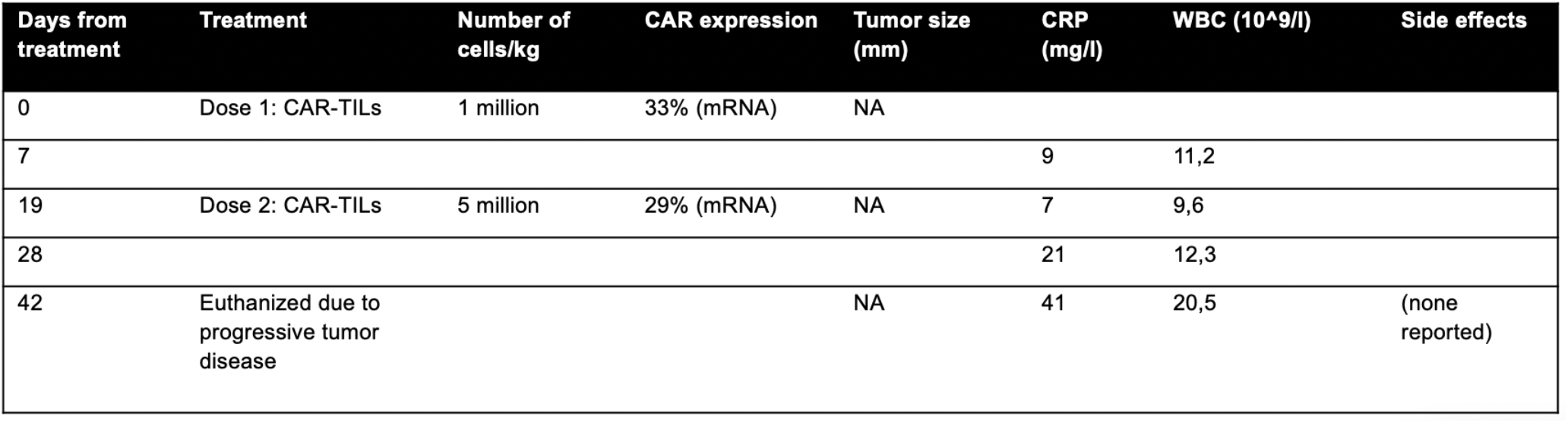
Clinical responses to CAR-TIL treatment of Dog 2 (Bella)

| Days from treatment | Treatment | Number of cells/kg | CAR expression | Tumor size (mm) | CRP (mg/l) | WBC (10<sup>9</sup>/l) | Side effects |
| --- | --- | --- | --- | --- | --- | --- | --- |
| 0 | Dose 1: CAR-TiLs | 1 million | 33% (mRNA) | NA |  |  |  |
| 7 |  |  |  |  | 9 | 11,2 |  |
| 19 | Dose 2: CAR-TiLs | 5 million | 29% (mRNA) | NA | 7 | 9,6 |  |
| 28 |  |  |  |  | 21 | 12,3 |  |
| 42 | Euthanized due to progressive tumor disease |  |  | NA | 41 | 20,5 | (none reported) |

## Dataset

**Supplemental Table 3. Difference in expression of homing receptors in blood derived and tumor derived T-cells**

Statistical details for the differential expression analysis of single-cell RNA sequencing data underlying **Fig. S1a**.

